# Closed genomes uncover a saltwater species of *Candidatus* Electronema and shed new light on the boundary between marine and freshwater cable bacteria

**DOI:** 10.1101/2022.10.26.513921

**Authors:** Mantas Sereika, Francesca Petriglieri, Thomas Bygh Nymann Jensen, Artur Sannikov, Morten Hoppe, Per Halkjær Nielsen, Ian P.G. Marshall, Andreas Schramm, Mads Albertsen

## Abstract

Cable bacteria of the *Desulfobulbaceae* family are centimeter-long filamentous bacteria, which are capable of conducting long-distance electron transfer. Currently, all cable bacteria are classified into two candidate genera: *Candidatus* Electronema, typically found in freshwater environments, and *Candidatus* Electrothrix, typically found in saltwater environments. This taxonomic framework is based on both 16S rRNA gene sequences and metagenome-assembled genome (MAG) phylogenies. However, most of the currently available MAGs are highly fragmented, incomplete, and thus likely miss key genes essential for deciphering the physiology of cable bacteria. To address this, we performed Nanopore long read (total 162.4 Gbp) and Illumina short read (total 148.3 Gbp) shotgun sequencing of selected environmental samples and a single-strain enrichment of *Ca*. Electronema aureum. We recovered multiple cable bacteria MAGs, including two circular and one single-contig. Phylogenomic analysis, also confirmed by 16S rRNA gene-based phylogeny, classified one circular MAG and the single-contig MAG as novel species of cable bacteria, which we propose to name *Ca*. Electronema halotolerans and *Ca*. Electrothrix laxa, respectively. The *Ca*. Electronema halotolerans, despite belonging to the previously recognized freshwater genus of cable bacteria, was retrieved from brackish-water sediment. Metabolic predictions showed several adaptations to a high salinity environment, similar to the “saltwater” *Ca*. Electrothrix species, indicating how *Ca*. Electronema halotolerans may be the evolutionary link between marine and freshwater cable bacteria lineages.

## Introduction

Cable bacteria of the *Desulfobulbaceae* family (*Desulfobacterota*) are centimeter-long, multicellular filamentous bacteria capable of long-distance electron transfer^1–3^. According to the current taxonomic framework, they belong to genus *Candidatus* Electronema (freshwater-based) or *Candidatus* Electrothrix (saltwater-based)^4^. Cable bacteria can be found globally in freshwater and saltwater sediments^5–7^ as well as around oxygen-releasing roots of aquatic plants^8,9^.

The ecological borderline between saltwater and freshwater habitats continues to be an unresolved challenge in microbiology^10^. Even though these aquatic environments share some ecological features, the radical shift in salinity and ionic concentration suggests that the transition from high to low salinity habitats must be accompanied by substantial changes in the metabolic repertoire and cellular complexes, in response to physico-chemical shifts and substrate availability^10,11^. These mechanisms have not yet been clarified in cable bacteria, even though the Na^+^/H^+^ antiporter NhaA has been previously suggested as a potential discriminant between marine and freshwater cable bacteria^12^.

The current metabolic model proposes that cells belonging to one cable filament can exhibit two different kinds of physiological features: at the anoxic zone within the deeper layers of the sediment, cells oxidize sulfide and the resulting electrons are transferred along the conductive structure to the oxic zone, where the cells utilize them to reduce oxygen^12,13^. This long-distance electron transport (LDET) has been demonstrated using Raman microscopy^3^ and the conductive structure involved has recently been proposed to consist of carbohydrates and proteins containing a sulfur-ligated nickel group, which would be an unprecedented form of biological electron transport^14^.

Despite enrichment efforts^12,15^, cable bacteria species have not been isolated in pure culture. Therefore, the currently available approach for acquiring cable bacteria genomic sequences is through shotgun sequencing of microbial communities and recovery of metagenome-assembled genomes (MAGs). Although MAGs of cable bacteria have been published by previous studies^12,13,15^, the publically available cable bacteria genomes are presently highly fragmented, with multiple draft genomes missing 16S rRNA genes and exhibiting reduced genome completeness.

The main cause of existing cable bacteria MAGs featuring significant fragmentation and overall lower genome quality is due to using short read sequencing for genome-centric metagenomics, which often results in highly fragmented assemblies^16,17^, usually due to the reads not being able to span genomic repeats, resulting in contig breaks during assembly^18^. Binning of short contigs can also lead to incomplete genomes^19–21^ or genome bins contaminated with contigs from other organisms^22–24^. Long read sequencing (e.g. Oxford Nanopore or Pacific Biosciences) has been demonstrated by multiple studies to improve on metagenome assembly contiguity, resulting in less fragmented genome bins, as well as enabling the recovery of complete, circular bacterial genomes from complex samples^25–33^.

Here, we used deep Nanopore long-read metagenomic sequencing to recover closed and high-quality genome drafts of cable bacteria. Through metabolic annotation of the recovered MAGs and visualization of new cable bacteria species, we provide novel insights into the functional potential, morphology and ecological niches of these enigmatic bacteria.

## Results and Discussion

### Recovery and phylogenetic analysis of cable bacteria MAGs

One *Ca*. Electronema aureum GS enrichment (ENR) and two environmental (brackish-water sediment — BRK; marine sediment — MAR) samples, containing complex microbial communities with known cable bacteria populations^12,34^, were sequenced at varying depth (**Table S1**). In total, 162.4 Gbp and 148.3 Gbp of Nanopore and Illumina read data, respectively, was generated and yielded 103 high-quality (HQ) and 195 medium-quality (MQ) MAGs after automated binning. Overall, HQ and MQ MAGs accounted for approx. 40% of the cumulative relative abundance in long and short read datasets (per sample), with the MAR sample featuring the lowest-yielding automated binning metrics, while the enrichment sample exhibited the most contiguous metagenome assembly statistics, yielding 66 closed bacterial and archaeal MAGs. After manual inspection and reassembly, a total of 5 cable bacteria MAGs were recovered (**Table S2**), spanning both candidate genera of cable bacteria (**Figure 1**).

**Figure 1.**
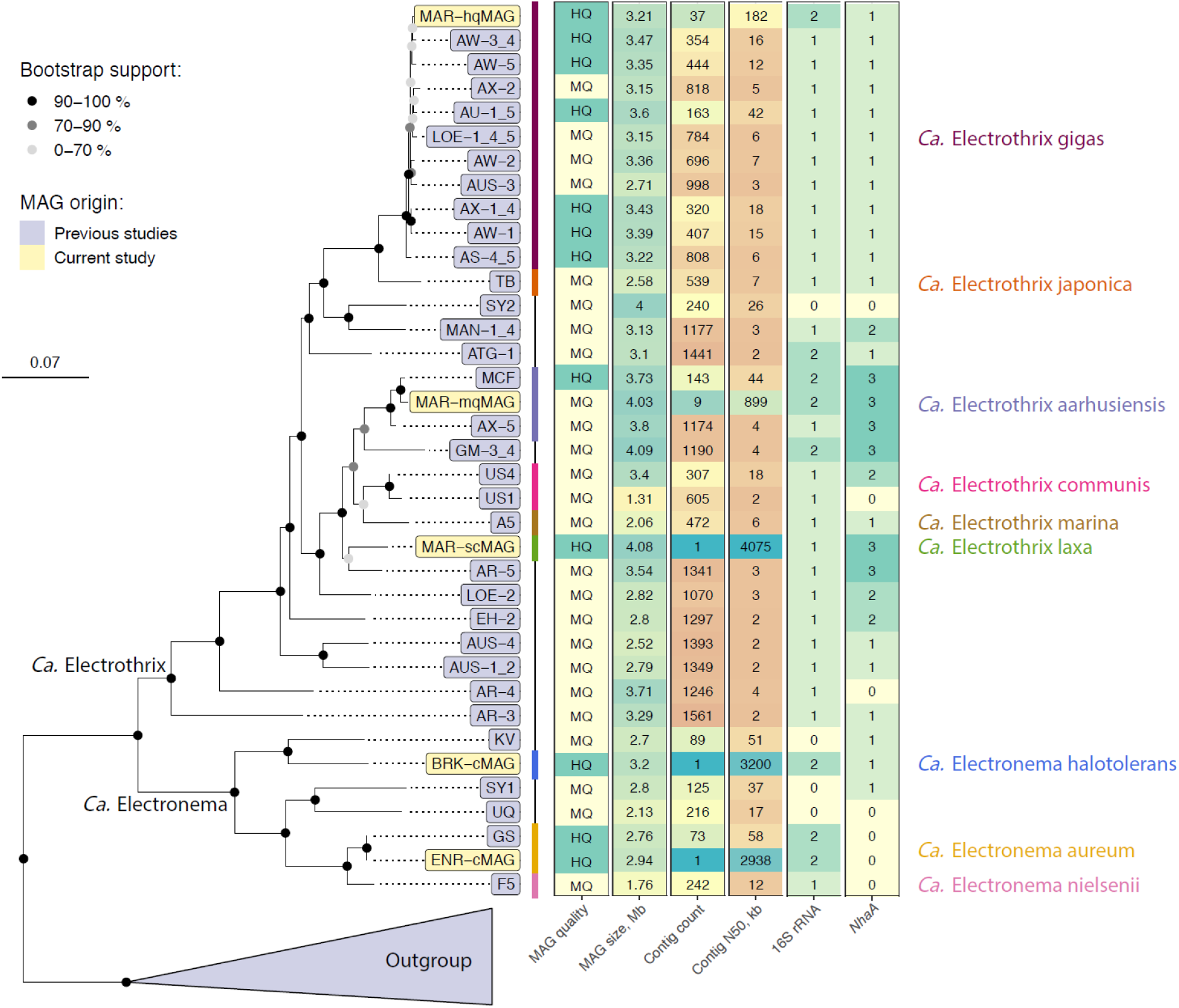
Cable bacteria genome tree and selected MAG features. The maximum-likelihood tree was built from 120 universal bacterial marker genes, 100 bootstraps, and multiple outgroups were included (**Table S5**). Only HQ and MQ MAGs were used for building the tree. Nodes suspected of delineating relevant phylogenetic groups are highlighted. MAG quality — quality ranking according to MIMAG standards. MAG size, Mb — total MAG size in megabases. Contig count — number of contigs per MAG. Contig N50, kb — MAG N50 values in kilobases. 16S rRNA — count of 16S rRNA gene sequences detected in MAGs. *NhaA* — count of *NhaA* gene sequences predicted in MAGs.

The ENR sample yielded a first-ever circular MAG (ENR-cMAG) of an enriched *Ca*. Electronema aureum GS strain, described in a previous study^12^. The circular genome featured < 0.1% SNP rate, suggesting minimal presence of strain microdiversity^35^. Compared to the previously published short-read MAG of the same strain, the ENR-cMAG featured 100% average nucleotide identity (ANI, **Figure S1**, **S2**) and identical 16S rRNA gene sequences (**Figure S3**), even though the closed MAG for *Ca*. Electronema aureum GS was found to contain more genes as well as more insertion sequence elements than the short-read-based equivalent MAG (**Table S3**). Furthermore, the previously published and fragmented *Ca*. Electronema aureum MAG had 4 contigs (encoding 47 genes) that did not align to the closed ENR-cMAG, which indicates minor contamination.

The MAR sample was found to contain genomic sequences of multiple cable bacteria and 3 MAGs were recovered of varying contiguity and quality: a single-contig, HQ MAG (MAR-scMAG), a HQ MAG of 37 contigs (MAR-hqMAG) as well as a MQ MAG of 9 contigs (MAR-mqMAG). All cable bacteria MAGs from the MAR sample featured above 90% genome completeness and higher than > 0.5% SNP rate, indicating the presence of some strain heterogeneity. Using ANI of 95% for species-level demarcation^36^, the MAR-mqMAG and MAR-hqMAG were found to be members of *Ca*. Electrothrix aarhusiensis and *Ca*. Electrothrix gigas^34^, respectively (**Figure S1**). Species-level assignments for MAR-mqMAG as well as MAR-hqMAG were further confirmed with 16S rRNA phylogeny (**Figure S3**) and the MAR-hqMAG was also observed to supplement the *Ca*. Electrothrix gigas species pangenome with more genes (**Table S4**).

More interestingly, the MAR-scMAG was found to be a member of *Ca*. Electrothrix (**Figure 1**), although the MAG featured less than 90% ANI to all existing HQ and MQ cable bacteria MAGs (**Figure S1**). The 16S rRNA tree further confirmed MAR-scMAG as being phylogenetically distinct from other *Ca*. Electrothrix 16S rRNA gene sequences. Thus, we suggest that MAR-scMAG belongs to a novel species in *Ca*. Electrothrix, which we have proposed to name *Ca*. Electrothrix laxa due to the large filament size of the species (see later).

The third sample sequenced in this study, BRK, yielded another circular cable bacterium MAG (BRK-cMAG) with < 0.1% SNP rate (**Table S2**). Surprisingly, the MAG was phylogenetically classified as *Ca*. Electronema using the genome tree (**Figure 1**) as well as 16S rRNA gene phylogeny (**Figure S3**), even though the sample was collected from a brackish-water sediment, where *Ca*. Electrothrix was expected to be present^37^. Using 75-77% ANI and 50% percentage of conserved proteins (POCP) for genus boundary^36,38,39^, the BRK-cMAG belongs to the *Ca*. Electronema genus (**Figure S1-2**). While just inside the established ANI and POCP genus boundaries, the BRK-cMAG had less than 78% ANI to all cable bacteria MAGs (**Figure S1**). Clustering all cable bacteria MAG genes at different identity cutoffs revealed that BRK-cMAG featured the highest fraction of unique genes, indicating that the BRK-cMAG exhibited the most novel gene content of the cable bacteria MAGs recovered in this study (**Figure S4**). Based on its ability to survive in a saline environment, we suggest to name the BRK-cMAG as *Ca*. Electronema halotolerans. Interestingly, undefined *Ca*. Electronema species have been previously detected by 16S rRNA gene amplicon sequences in root and associated bulk samples of the seagrass species *Halophila ovalis* and *Zostera marina*^9^. Therefore, we compared these undefined *Ca*. Electronema ASVs with the 16S rRNA gene sequence of *Ca*. Electronema halotolerans. The phylogenetic analysis (**Figure S3**) confirmed the association of such ASVs with the new *Ca*. Electronema species and further indicated its potential tolerance of higher salinity.

### Adaptation to marine environment

Typically, halophilic microorganisms can use two different strategies to maintain osmotic balance inside their cells: the first, known as salt-in strategy, involves the intracellular accumulation of high concentration of salts, in particular potassium and chloride; the second, also known as the “compatible solute” strategy, involves the intracellular accumulation of organic compounds like polyols, betaines or ectoines, which can be synthesized by the cells themselves or taken up from the environment^40^. Therefore, we screened the cable bacteria genomes for potential adaptations to high salinity habitats, including mechanisms to maintain a high osmotic pressure inside the cells, tolerance to heavy metal ions or accumulation of osmotic solutes^41,42^. Ion pumps are usually involved in achieving ion gradients across the cell membrane, in particular to exclude Na^+^ and accumulate K^+^ and Cl^−42^. One of the most common is the Na^+^/H^+^ antiporter *NhaA*, which is detected in the genetic potential of all *Ca*. Electrothrix genomes and in *Ca*. Electronema halotolerans, and has been previously suggested as a potential discriminant between marine and freshwater cable bacteria^12^ (**Figure 2–3**, **Dataset S1**). The medium quality MAGs KV^43^ and SY1^44^, which are closely related to *Ca*. Electronema halotolerans, encoded also the *NhaA* gene (**Figure 2**), confirming its importance for survival of bacteria in saltwater environment^45^.

**Figure 2.**
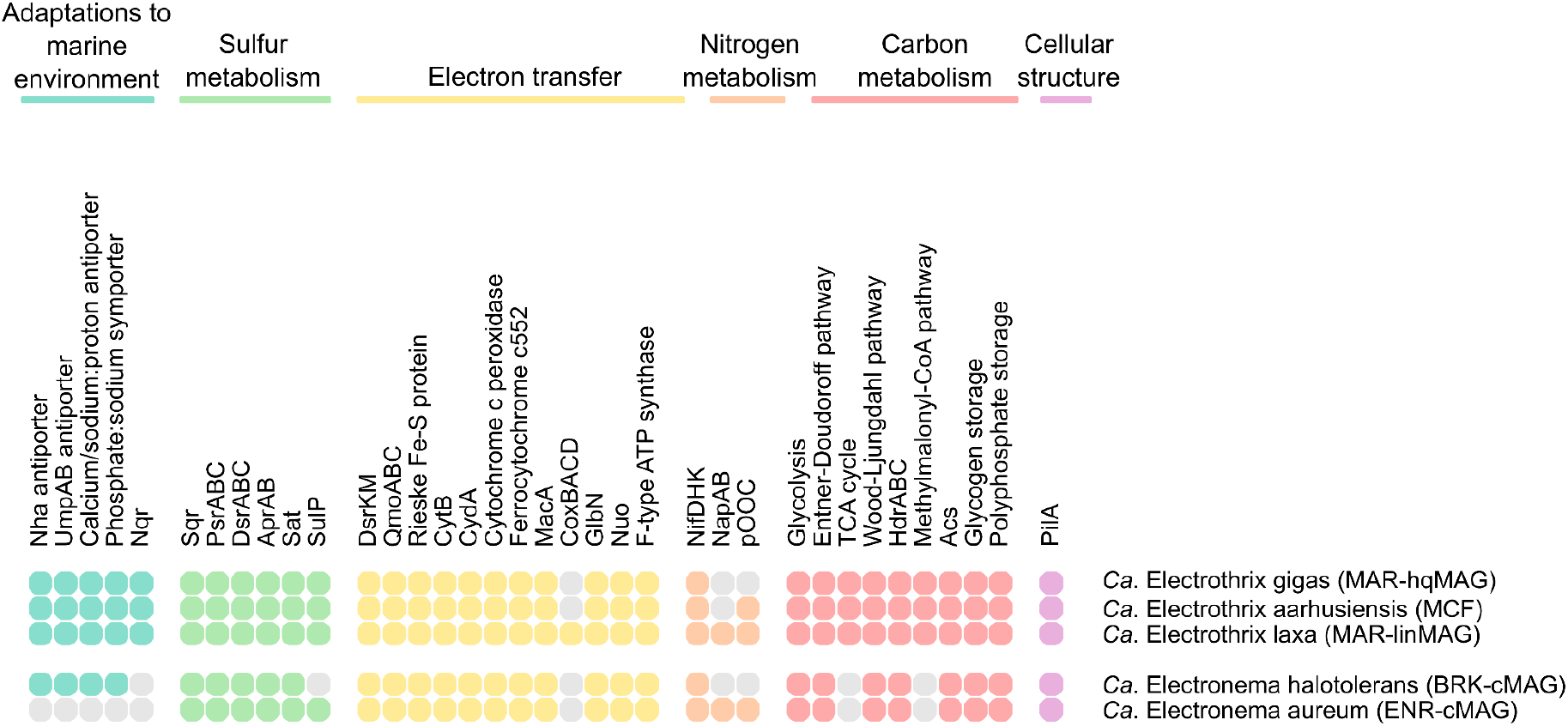
General overview of the functional potential of HQ cable bacteria MAGs. Pathways are considered present if more than 80% of the genes are predicted. For the full list of gene names and associated KO numbers see **Dataset S1**. The MAGs and genomes are ordered as in the genome tree in Figure 1.

**Figure 3.**
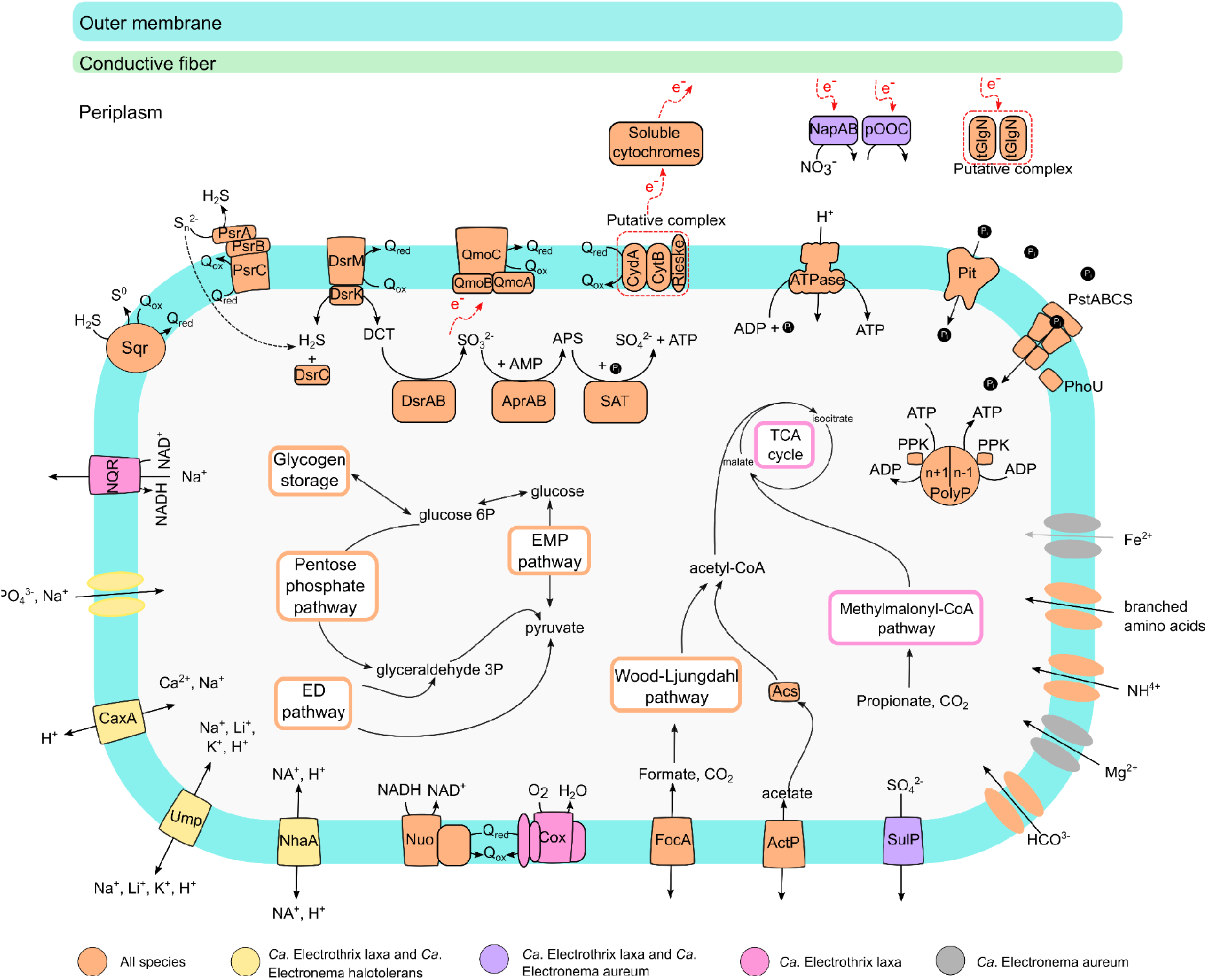
Metabolic model for *Ca*. Electronema halotolerans and *Ca*. Electrothrix laxa. Gray and red dotted lines indicate putative reactions and complexes. Abbreviations: EMP pathway, Embden–Meyerhof–Parnas pathway (glycolysis); TCA cycle, tricarboxylic acid cycle; EMP pathway, Entner-Doudoroff pathway; Acs, acetyl-CoA synthetase; Dsr, dissimilatory bisulfite reductase; DCT, DsrC-trisulfide; Apr, Adenosine phosphosulfate reductase; Sat, ATP sulfurylase; Psr, polysulfide reductase; SQR, sulfide:quinone oxidoreductase; Qmo, quinone-interacting membrane-bound oxidoreductase complex; Q(ox/red), quinone (oxidized or reduced); CydA, membrane-bound cytochrome bd quinol oxidase–subunit A; CytB, cytochrome bc complex–subunit B; Rieske, Rieske Fe-S domain protein; Nap, periplasmic nitrate reductase; pOOC, periplasmic multiheme cytochrome (nitrite reductase); tGlnN, truncated hemoglobin; PolyP, polyphosphate; PpK, polyphosphate kinase; Pit, inorganic phosphate transporter family; PstABCS, inorganic phosphate ABC transporter, PhoU, phosphate transport system accessory protein; SulP, sulfate permease; ActP, acetate transporter; FocA, putative formate transporter; Cox, cytochrome c oxidase; Nuo, NADH dehydrogenase; NhaA, Sodium:proton antiporter; Ump, putative sodium, lithium, potassium: proton antiporter; Cax, putative calcium, sodium:proton antiporter; NQR, sodium-translocating NADH:ubiquinone oxidoreductase. Further details are provided in the main text and supplementary notes.

We found three distinct clades of the gene encoding the NhaA Na^+^/H^+^ antiporter in *Ca*. Electronema and *Ca*. Electrothrix (**Figure S5a**), which appear to have been present in the common ancestor to the two genera. *NhaA_1* persisted in all *Ca*. Electrothrix, *NhaA_2* in *Ca*. Electrothrix aarhusensis and *Ca*. Electrothrix laxa, and *NhaA_3* in *Ca*. Electronema (**Figure S5b**). The high identity between the *NhaA_3* sequence in *Ca*. Electrothrix aarhusensis and *Ca*. Electrothrix laxa suggests that there has been a horizontal gene transfer event between *Ca*. Electrothrix laxa and *Ca*. Electrothrix aarhusensis. Furthermore, *Ca*. Electronema aureum lost all copies of *NhaA*, which likely contribute to the restriction of its habitat to freshwater. *NhaA* has been shown to confer saltwater tolerance when overexpressed in a freshwater cyanobacterium in the past^46^ and it appears that the presence of this gene is critical for the survival of *Ca*. Electronema in saltwater. It is unclear what the functional difference between the three different types of *NhaA*, but it is worth noting that all saltwater *Ca*. Electronema genomes found so far have been found in brackish water (salinity <30), suggesting that *NhaA_3* may be adapted to lower Na^+^ concentrations.

Interestingly, all *Ca*. Electrothrix laxa, *Ca*. Electrothrix gigas and *Ca*. Electronema halotolerans MAGs also encoded homologs (56% amino acid identity) of the two subunits of the Na^+^/Li^+^/K^+^:H^+^ antiporter UmpAB, a transporter firstly characterized in *Halomonas zhaodongensis^47^* (**Figure 2–3**, **Dataset S1**). Homologs of a putative Ca^2+^/Na^+^:H^+^ antiporter (41% amino acids identity with a transporter first identified in *Alkalimonas amylolytica*^48^) and a putative phosphate:Na^+^ symporter were present in *Ca*. Electrothrix and *Ca*. Electronema halotolerans MAGs, but absent in all the other *Ca*. Electronema genomes (**Figure 2–3**, **Dataset S1**).

The detection of several genes involved in ion translocation through the membrane may indicate the preference of the marine/brackish cable bacteria for a “salt-in” strategy^42^. This seems further supported by the fact that none of the cable bacteria MAGs encode the biosynthetic pathways for osmotics solutes and only one subunit of a putative glycine betaine/carnitine/choline transporter is detected in some of the MAGs, including the freshwater *Ca*. Electronema (**Dataset S1**).

### Closed genomes clarify key features of the cable bacteria metabolism

Metabolic prediction included a total of 10 HQ MAGs, while the remaining low and middle quality genomes were used as additional information to support the proposed metabolic model. All the MAGs retrieved in this study presented a metabolic potential consistent with the previously proposed model for *Ca*. Electronema and *Ca*. Electrothrix^12^ (**Figure 2–3**, **Dataset S1**). Furthermore, the availability of closed genomes allowed us to shed light on metabolic features of cable bacteria that were previously uncertain. One of the most controversial features of the long-distance electron transport (LDET) is the lack in cable bacteria genomes of a terminal oxidase, indicating the reduction of terminal acceptors without energy conservation^12^. This is confirmed by our findings, with no known complete terminal cytochrome bd-II oxidase detected in the closed genomes and other MAGs. However, the MAGs encoded only homologs of the subunit CydA as previously reported^12^ (**Figure 2–3**, **Dataset S1**). Interestingly, the genome of *Ca*. Electrothrix laxa encodes all four subunits of the membrane-bound cytochrome c oxidase (Cco), with ~60% amino acid sequence identity to homologs in other *Desulfobulbaceae* species and most likely resulting of horizontal gene transfer, as previously observed for its close relative *Ca*. Electrothrix communis^12^ (**Figure 2–3**, **Dataset S1**). We can therefore hypothesize that these species may be able to couple oxygen reduction to proton translocation with energy conservation.

All cable bacteria show the metabolic potential for nitrogen fixation (**Figure 2–3**, **Dataset S1**) via the molybdenum-dependent nitrogenase NifDHK. The ammonium, which can also be taken up by the AmtB ammonium transporter, can then be incorporated into amino acids via glutamine/glutamate synthesis (**Dataset S1**). Experimental evidence has shown that cable bacteria can couple sulfide oxidation with nitrate or nitrite reduction^13,49^. Only two species (*Ca*. Electronema aureum and *Ca*. Electrothrix laxa) have the potential for nitrate reduction via the periplasmic nitrate reductase NapAB, and nitrate reduction by the NapAB system has been recently confirmed by transcriptomics in *Ca*. Electronema aureum^49^. The *nap* operon is lacking the gene encoding for the subunit NapB in *Ca*. Electrothrix laxa, but NapB has been shown to be non-essential in the nitrate reduction of *Shewanella oneidensis*^50^. No Nir- or Nrf-type nitrite reductase were detected in *Ca*. Electronema aureum and *Ca*. Electrothrix laxa (**Figure 2–3**, **Dataset S1**), but a periplasmic multiheme cytochrome adjacent to the nap operon, which is also predicted in *Ca*. Electrothrix aarhusiensis, has been suggested as the enzyme responsible for nitrite reduction to ammonium^12,49^, suggesting a linked function and preference of cable bacteria for dissimilatory nitrate reduction to ammonium (DNRA).

All the MAGs encoded nearly complete central carbon processing pathways through glycolysis, Entner–Doudoroff pathway, pentose phosphate pathway and TCA cycle (**Figure 2–3**, **Dataset S1**). As previously reported for *Ca*. Electronema aureum and *Ca*. Electrothrix aarhusiensis, and now confirmed by the availability of closed genomes, no genes encoding for the enzyme enolase were detected in the MAGs, indicating a potential alternative and unknown pathway to complete glycolysis^12^. In the carbon metabolism it’s residing also one of the major differences in the metabolic potential of the two genera of cable bacteria. While *Ca*. Electrothrix MAGs encoded the potential for full TCA cycle, the two closed genomes for *Ca*. Electronema MAGs lacked several genes, including succinyl-CoA-synthase 2-oxoglutarate ferredoxin oxidoreductase (**Figure 2–3**, **Dataset S1**). Similarly, all *Ca*. Electrothrix MAGs have the potential to assimilate propionate via the methylmalonyl-CoA pathway to succinyl-CoA, absent in *Ca*. Electronema (**Figure 2–3**, **Dataset S1**).

### Visualization of the new species

A set of new FISH probes (**Table S6**) was designed to distinguish the species *in situ* (**Figure 4**). The diameter of filaments ranged from 1 to 6 μm for *Ca*. Electrothrix laxa and 1 to 2 μm for *Ca*. Electronema halotolerans. Raman microspectroscopy has recently proved to be an excellent technique in helping to clarify the structure of the conductive structure of cable bacteria^14^. Therefore, we applied it in combination with the newly designed FISH probes to confirm the chemical composition of the cable’s cell envelope in the new species. Raman spectra of FISH-identified *Ca*. Electrothrix laxa showed peaks typical of biological components, such as phenylalanine (1005 cm^−1^), CH_2_ bond (1462 cm^−1^) and the amide I peak of protein (1672 cm^−1^), as well as peaks linked to cytochromes (750 cm^−1^, 1129 cm^−1^, 1314 cm^−1^ and 1586 cm^−1^). Additionally, two large peaks were present at 371 and 492 cm^−1^ (**Figure S6**), which may be linked to the presence of a S-ligated metal group in the conductive fiber^14^.

**Figure 4.**
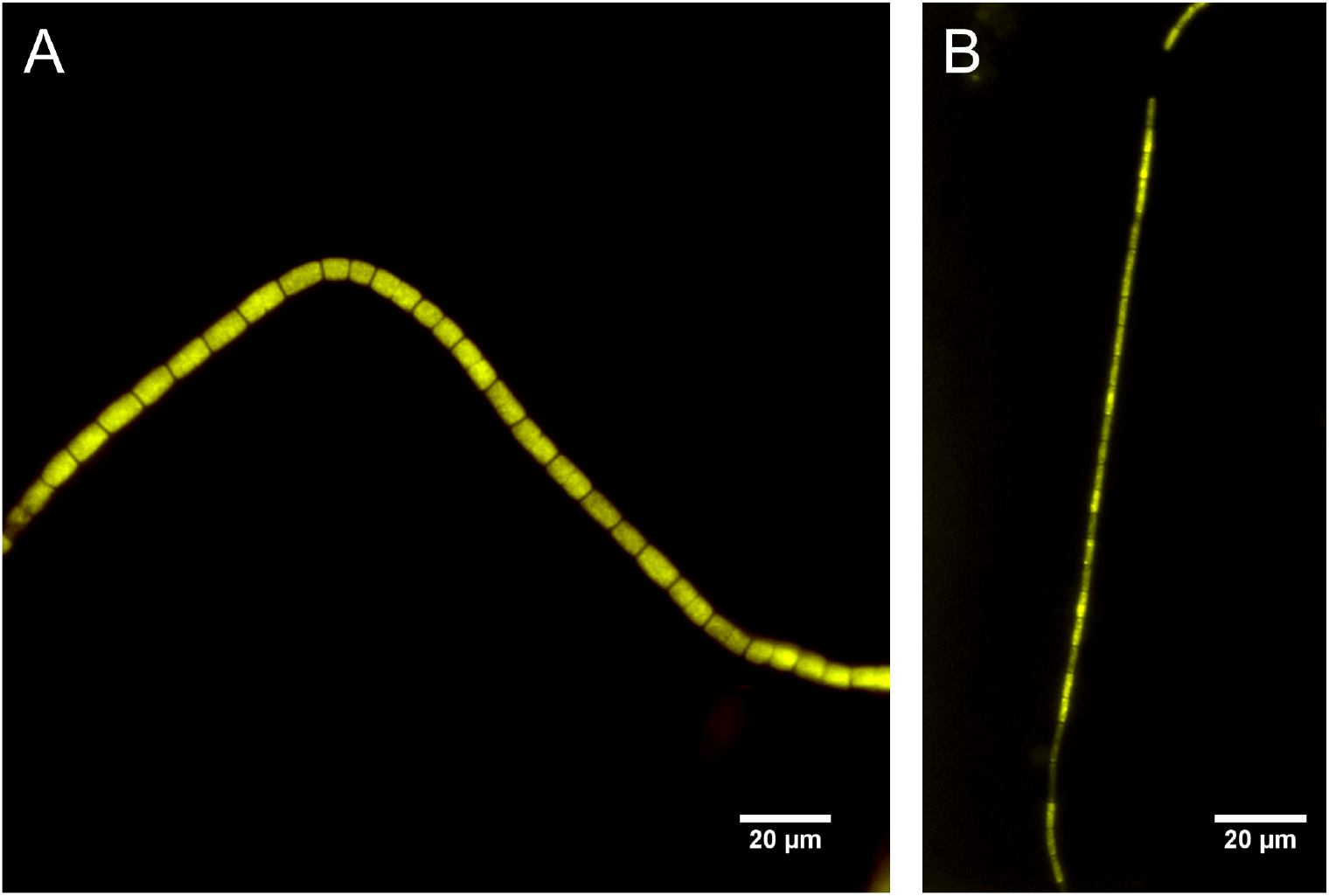
Composite FISH micrographs of the two novel cable bacteria species. **A)** *Ca*. Electrothrix laxa appears in yellow, by the overlap of the species-specific probe Ex-la-189 (FAM-labeled, green) and the probe for the *Desulfobulbaceae* family DSB706 (Cy3-labeled, red); **B)** *Ca*. Electronema halotolerans appears in yellow, by the overlap of the species-specific probe EN-ha-80 (FAM-label, green) and the probe for the *Desulfobulbaceae* family DSB706 (Cy3, red).

## Conclusion

With this study, we provide the first-ever closed genomes of cable bacteria. The closed genome for *Ca*. Electronema aureum enabled the identification of novel genes and genomic repeats, which were absent from the previously published fragmented MAGs, as well as confirm metabolic models proposed for cable bacteria. Furthermore, we also provide the first-ever single-contig HQ MAG for *Ca*. Electrothrix which we propose to name as novel species of *Ca*. Electrothrix laxa, which we suspect is capable of functions atypical for cable bacteria due to horizontal gene transfer. Lastly but most importantly, we provide a closed genome of a novel species we propose to name *Ca*. Electronema halotolerans, which is the first-ever genome draft for a *Ca*. Electronema that was observed to be present and feature genes for survival in a saltwater environment. The existence of a saline-tolerant *Ca*. Electronema species challenges the current freshwater-saltwater-centric taxonomic framework for cable bacteria, necessitating a re-assessment for the categorization of cable bacteria candidate genera.

### Taxonomic proposal

Description of *‘Candidatus* Electrothrix laxa’ sp. nov.: *“Candidatus* Electrothrix laxa*”*, (la’xa, L. fem. adj. *laxa*, large). This taxon is represented by the MAG MAR-scMAG. The complete protologue can be found in **Table S7**.

Description of *‘Candidatus* Electronema halotolerans’ sp. nov.: *“Candidatus* Electronema halotolerans”, (ha.lo.to’le.rans, from Gr. n. *hals*, halos salt; L. part. adj. *tolerans* tolerating; N.L. part. adj. halotolerans). This taxon is represented by the MAG BRJ-cMAG. The complete protologue can be found in **Table S8**.

## Methods

### Sample collection and manual processing

A marine sediment sample was collected from Hou, Denmark (55°54’36.8”N 10°14’46.8”E) and the brackish-water sediment sample was collected from Løgten, Denmark (56°17’17.8”N 10°22’54.9”E). *Ca*. Electronema aureum GS enrichment sample and manual processing of all samples was performed as detailed in a previous study^15^.

### DNA Extraction and QC

DNA from environmental samples was extracted using the DNeasy PowerSoil Pro Kit (QIAGEN, Germany) according to the manufacturer’s protocol. The concentration of the extracts was measured using the Qubit dsDNA HS kit (Thermo Fisher Scientific, USA, #Q33231) with a Qubit 3.0 fluorometer (Thermo Fisher Scientific, USA), while DNA purity was assessed with a NanoDrop One Spectrophotometer (Thermo Fisher, USA). Sample DNA fragment sizes were inspected using an Agilent 2200 Tapestation system with Genomic DNA ScreenTapes (Agilent Technologies, USA, #5067-5365). DNA samples were also size-selected (with the exception of BRK sample due to limited DNA amounts) using the Circulomics SRE XS kit (Circulomics, USA), following the manufacturer’s instructions to deplete DNA fragments below 10 kb.

### Nanopore DNA sequencing

DNA libraries for Nanopore sequencing were prepared using the SQK-LSK110 Ligation Sequencing kit (Oxford Nanopore Technologies, UK) according to the manufacturer’s instructions. The libraries were loaded into Nanopore R.9.4.1 chemistry flow cells and sequenced on the MinION Mk1B sequencing device (Oxford Nanopore Technologies, UK) using the MinKNOW v21.06.13 (https://community.nanoporetech.com/downloads) software. Raw Nanopore read data was basecalled with Guppy (v. 5.0.7, https://community.nanoporetech.com/downloads) using the “dna_r9.4.1_450bps_sup.cfg” model.

### Illumina DNA sequencing

Short read sequencing libraries were prepared using the Illumina DNA Prep kit in combination with IDT^®^ UD Indexes Set A (Illumina, USA, #20018705, #20027213). Between 66.3 ng and 200 ng per sample was used as starting material. 5 cycles of PCR was applied in the amplification of the tagmented DNA. All other steps were performed according to the manufacturer’s protocol. Final libraries were quantified and insert size evaluated using the Qubit dsDNA HS assay (Thermo Fisher Scientific, USA, #Q33231) and a D1000 screentape (Agilent Technologies, USA, #5067-5582, #5067-5602). Samples were multiplexed using 100 ng of DNA and the final pooled library was evaluated in the same way as for the individual libraries.

The pooled library was sequenced on an S1 flow cell for the NovaSeq 6000 platform (Illumina, USA). 2.0 nM was used as input for the V1.5 300 cycle kit (Illumina, USA, #20012863).

### Sequencing read processing

Illumina reads were trimmed, quality-filtered and de-duplicated using fastp (v. 0.23.2^51^) with “-l 100 --dedup --dup_calc_accuracy 6” settings. For Nanopore reads, adaptor sequences were trimmed off and chimeric reads were split using Porechop (v. 0.2.3^52^). Reads with a lower Phred score than 7 or shorter read length than 200 bp were discarded using NanoFilt (v. 2.6.0^53^). Read length and quality statistics were acquired with NanoPlot (v. 1.24.0^53^).

### Metagenome assembly, polishing and binning

Trimmed and filtered nanopore reads were assembled into metagenomes via the Flye (v. 2.9-b1768^54^) assembler using the “--meta”, “--nano-hq” and “min_read_cov_cutoff=8” settings. The assembly was then polished first with Nanopore reads using 3 rounds of Racon (v. 1.3.3^55^) and then 2 rounds of Medaka (v. 1.4.4, https://github.com/nanoporetech/medaka) polishing with “r941_min_sup_g507” model. The metagenome was then polished using Illumina reads with 1 round of Racon. Dependencies for metagenome polishing, read mappings include Minimap2 (v. 2.24^56^) and SAMtools (v. 1.14^57^). Contigs with lower length than 1kb were removed via Bioawk v. 1.0 (https://github.com/lh3/bioawk).

The assembled metagenomes were then binned using MetaBAT2 (v. 2.12.1^58^), with “-s 500000” setting, Vamb (v. 3.0.2^59^) with “-o C--minfasta 500000” setting, MaxBin2 (v. 2.2.7^60^) and metaBinner (v. 1.4.2^61^). Contig coverage profiles from Nanopore and Illumina read data were used as input for binning. The bins were then refined using DAS Tool (v. 1.1.2^62^) with “--search_engine diamond” setting. For the enrichment sample, all circular contigs longer than 500 kb were selected and used as bins. CoverM (v. 0.6.1, https://github.com/wwood/CoverM) was used to acquire bin coverage (“-m mean” setting) and relative abundance (“-m relative_abundance” setting) values.

### Bin processing and reassembly

Genome bin completeness and contamination values were acquired with CheckM (v. 1.1.2^23^) using the lineage-specific workflow and the Deltaproteobacteria marker lineage (UID3217) was used for all cable bacteria MAGs. Bin rRNA genes were predicted with Barrnap (v. 0.9, https://github.com/tseemann/barrnap) and tRNA gene sequences were predicted using tRNAscan-SE (v. 2.0.5^63^). Bin quality classification was performed in accordance with the Genomic Standards Consortium guidelines, where a high quality genome bin featured greater genome completeness than 90%, lower genome contamination than 5%, a minimum of 18 different tRNA genes, and the 16S, 5S, 23S rRNA genes occurring at least once in the bin^64^. Furthermore, bins with completeness greater than 50% and contamination lower than 10% were assigned as medium quality, whereas the remaining bins with less than 10% contamination were classified as low quality. Bins with contamination estimates greater than 10% were considered contaminated. Bins were de-replicated between the different samples using dRep (v. 2.6.2, https://github.com/MrOlm/drep) with “-comp 50 -con 10 -sa 0.95” settings.

Bin taxonomy was assigned using GTDB-Tk (v. 2.0.0^65^, R207 database) with the “--full_tree” setting enabled. Contig taxonomic classification was carried out using mmseqs2 (v. 13.45111^66^) with NCBI nr protein database^67^ (download date: 2021-06-26). 16S rRNA genes were classified using usearch (v. 11.0.667^68^) “-usearch_global” command against the SILVA v138 nr99 database^69^.

To acquire the circular cable bacteria genome from BRK sample, reads were extracted for contigs present in bins classified as cable bacteria by GTDB-Tk using SAMtools bam2fq command on the read mapping files and the extracted Nanopore reads were assembled with Flye. For the MAR sample, 2 reassemblies were performed. For the first one, which resulted in the linear cable bacteria genome, reads were extracted for contigs above 5 kb length, classified as *Desulfobulbaceae* by mmseqs2 or containing *Desulfobulbaceae* 16S rRNA genes or from MAGs classified as *Desulfobulbaceae* by GTDB-tk. For the second reassembly, which allowed to recover 2 fragmented cable bacteria MAGs from the MAR sample, a subset of reads from the first reassembly was made by extracting reads from contigs with longer length than 10 kb and higher coverage than 100, and after reassembly with Flye (using the “min_read_cov_cutoff=80” setting), the bins were manually refined by cross-referencing contigs with the Flye assembly graph. The acquired cable bacteria MAGs and sequenced reads are publicly available via accession numbers provided in **Table S7**.

### Comparative genomics and MAG annotation

For constructing cable bacteria genome trees, a multiple sequence alignment for 120 bacterial maker genes was generated using GTDB-Tk and provided to IQ-TREE (v. 2.0.6^70^) to build a tree using the “WAG+G” model and 100x bootstrapping. Multiple outgroups were included in the tree, summarized in **Table S5**. ANI and alignment coverage values between different cable bacteria MAGs and outgroups were acquired via pyANI (v. 0.2.11^71^) with “-m ANIb” setting enabled. AAI and POCP measurements between the genomes were acquired with CompareM (v. 0.1.2, https://github.com/dparks1134/CompareM) using the “--evalue 0.00001 --per_identity 40 --per_aln_len 50 --blastp” options. Quast (v. 4.6.3^72^) was used to compare MAGs of closely-related strains and ISEScan (v. 1.7.2.3^73^) was applied to identify insertion sequence elements.

MAGs were automatically annotated using Prokka (v. 1.14.0^74^) with “--metagenome” setting enabled. Protein sequences annotated as *NhaA* were extracted, aligned using MUSCLE (v. 5.0.1428^75^) with 1000 iterations, followed by protein tree generation using IQ-TREE with “LG+G4” model and 1000x bootstrapping. MAG gene clustering and pangenome creation was performed by using Prokka output files as input for Roary (v. 3.13.0^76^) with “-cd 85” setting and varying identity thresholds, which are described in the text. Additionally, HQ-MAGs were selected based on quality standards proposed by Bowers et al.^64^, (completeness and contamination of > 90% and < 5%, respectively) and uploaded to the ‘MicroScope Microbial Genome Annotation & Analysis Platform’ (MAGE)^77^ for manual inspection and metabolic pathways were predicted by KEGG^78^ pathway profiling of MAGE annotations.

### 16S rRNA gene-based phylogenetic analysis, FISH probe design and evaluation and Raman microspectroscopy

Phylogenetic analysis of 16S rRNA gene sequences and design of FISH probes were performed using the ARB software v.6.0.6^79^. A phylogenetic tree was calculated based on comparative analysis of aligned 16S rRNA gene sequences, retrieved from SILVA database^80^ using the maximum likelihood method (PhyML) and a 1000 – replicates bootstrap analysis. Coverage and specificity were evaluated and validated *in silico* with the MathFISH software for hybridization efficiencies of target and potentially weak non-target matches^81^. When needed, unlabelled competitors and helper probes were designed. All probes were purchased from Biomers (Ulm, Germany), labelled with 6-carboxyfluorescein (6-Fam) or indocarbocyanine (Cy3) fluorochromes.

FISH was performed as described by Daims et al.,^82^. Optimal formamide concentration for each novel FISH probe was visually determined by microscope observation after performing hybridization at different formamide concentrations in the range 15-70% (with 5% increments). Optimal hybridization conditions are described in **Table S6**. EUBmix^83,84^ was used to target all bacteria and NON-EUB^85^ was used as a negative control for sequence independent probe binding. Microscopic analysis was performed with Axioskop epifluorescence microscope (Carl Zeiss, Germany) equipped with a LEICA DFC7000 T CCD camera or a white light laser confocal microscope (Leica TCS SP8 X). Raman microspectroscopy was applied in combination with FISH as previously described^86^ to identify cellular components and clarify the structure of the conductive fiber.

## Data availability

Sequencing read data is available at ENA with bio project ID: PRJEB52550. Code and datasets used to generate the figures as well additional resources are available at https://github.com/Serka-M/Cable-Bacteria-MAGs. GTDB-tk database used in the project can be accessed at https://data.ace.uq.edu.au/public/gtdb/data/releases/release207/. NCBI nr protein database can be downloaded from https://ftp.ncbi.nlm.nih.gov/blast/db/FASTA/. The SILVA database can be accessed at https://www.arb-silva.de/download/arb-files.

## Acknowledgments

This study was funded by a research grant from Poul Due Jensen Foundation (Microflora Danica) and by the Danish National Research Foundation (DNRF136). We thank Jesper L. Wulff for help with the sampling.

## Author Contributions Statement

MA, PHN, IM and AS designed the study. IM and AS oversaw the manual processing of the samples. MS extracted the DNA and performed Nanopore sequencing, while TBNJ generated the Illumina data. MS performed bioinformatics processing, MAG generation and comparative genomics. FP performed genome annotation, phylogenetic analysis and FISH/RAMAN microscopy. MS and FP wrote the initial manuscript. All authors reviewed the manuscript.

## Competing Interests Statement

All authors declare no conflict of interest.

## Notes

### Competing Interest Statement

The authors have declared no competing interest.

