## Supplementary information for "Closed genomes uncover a saltwater species of *Candidatus* Electronema and shed new light on the boundary between marine and freshwater cable bacteria"

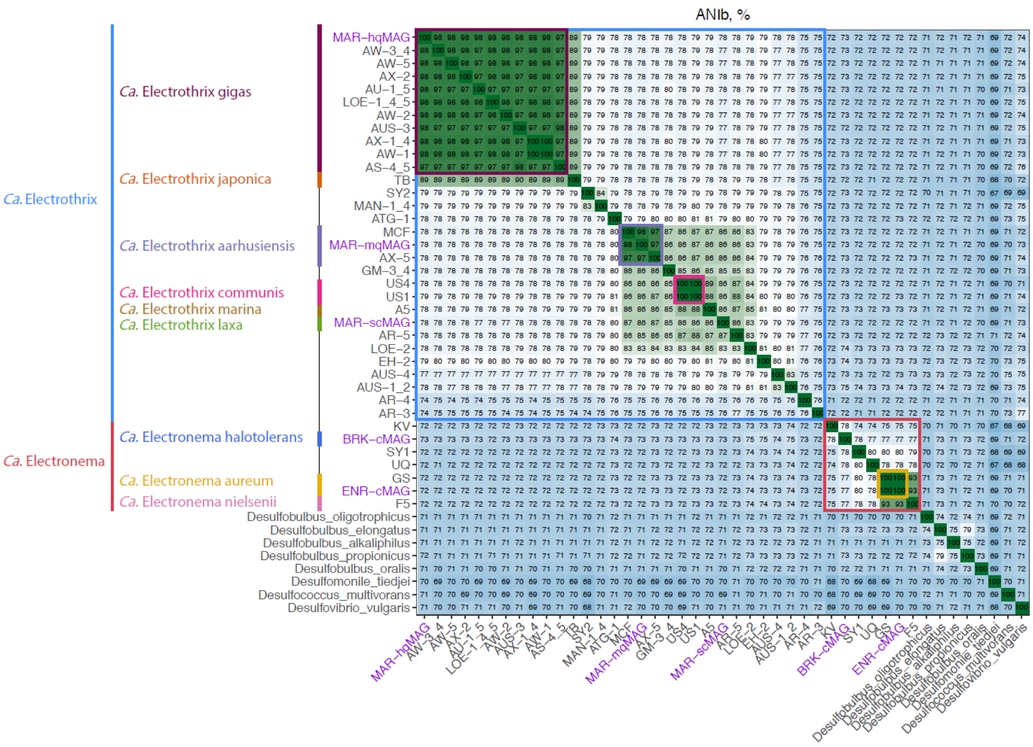


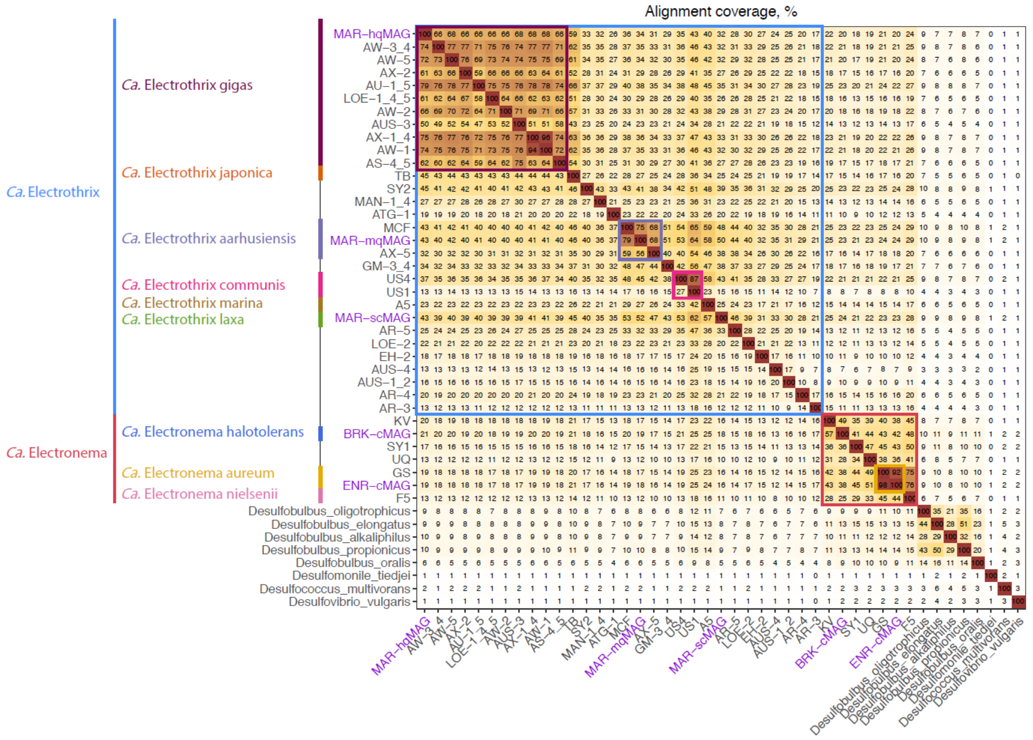


**Figure S1**. Genome comparison metrics for cable bacteria MAGs. ANI using BLAST (ANIb) values and genome alignment fraction between all selected MAGs are presented. MAGs recovered in this study as well as relevant phylogenetic groups are highlighted. Outgroups used in genome tree building were also included in the plots for context.

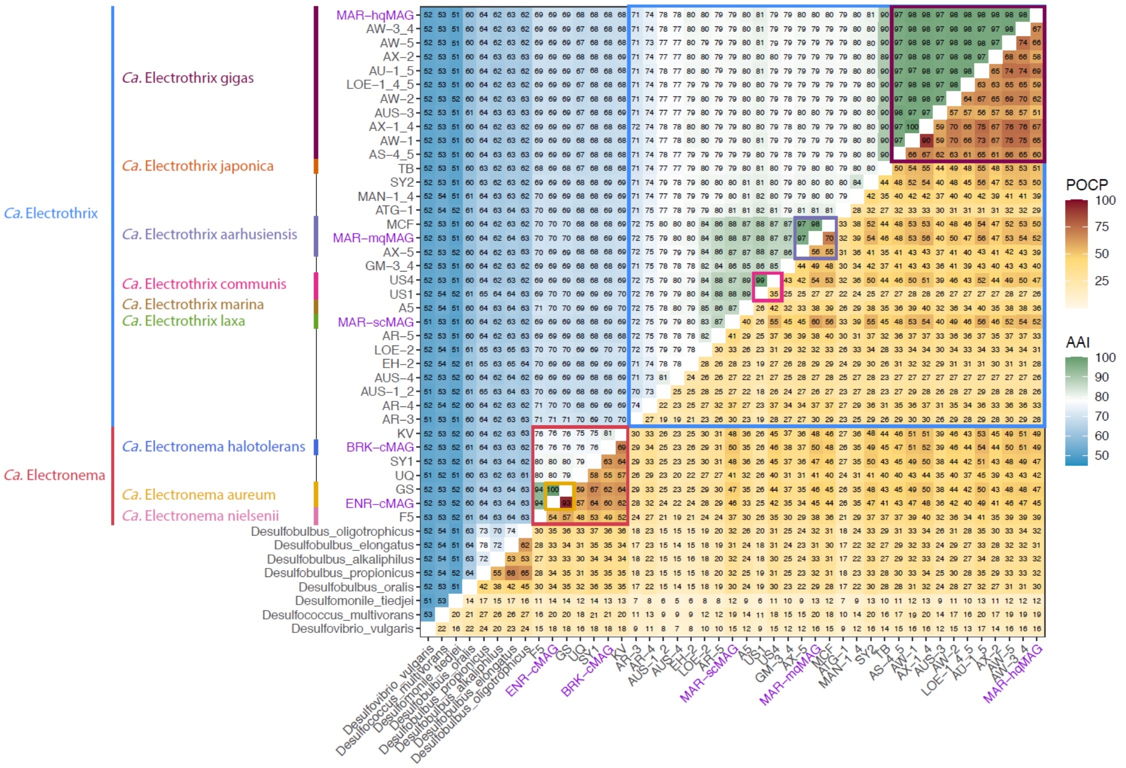


**Figure S2**. Protein comparison metrics for cable bacteria MAGs. AAI (top-left triangle) and POCP (bottom-right triangle) are presented. MAGs recovered in this study as well as relevant phylogenetic groups are highlighted. Outgroups used in genome tree building were also included in the plots for context.

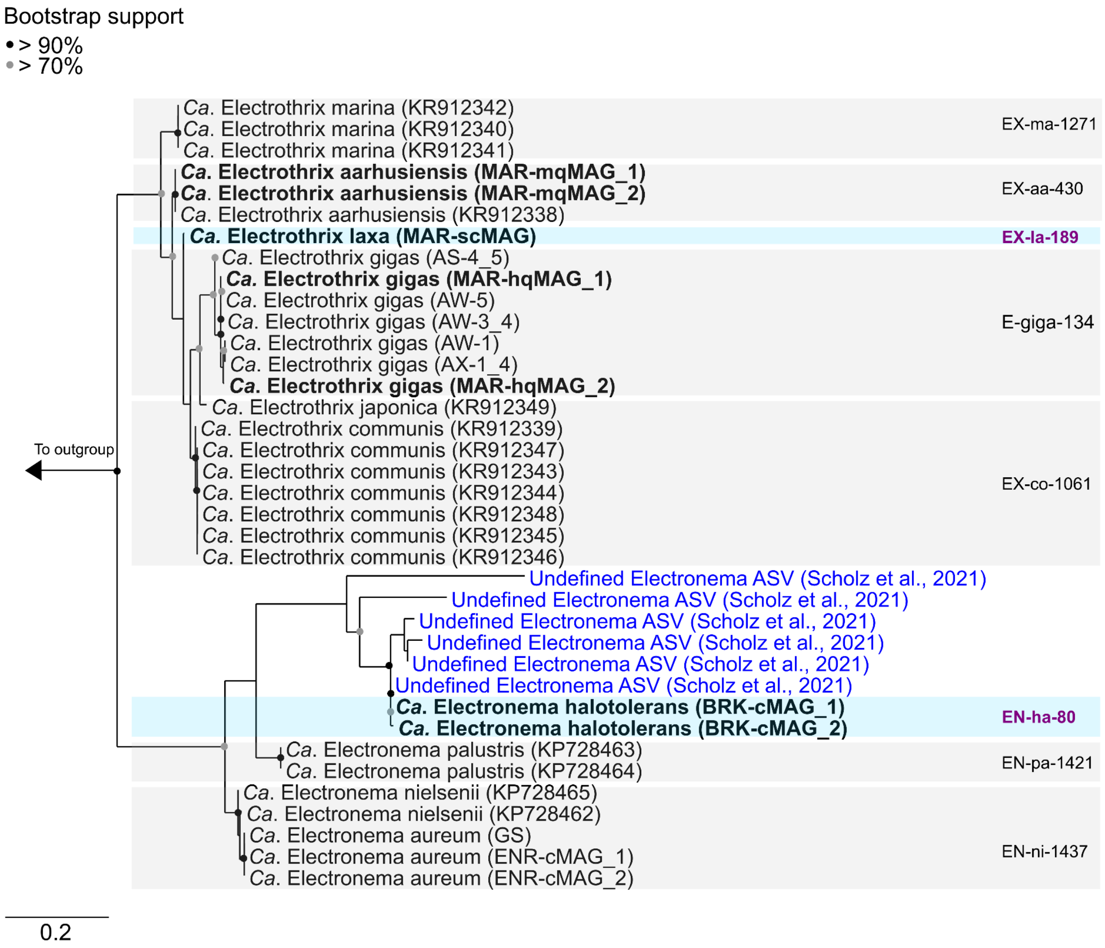


**Figure S3**. Maximum-likelihood (PhyML) 16S rRNA gene phylogenetic tree of known cable bacteria. 16S rRNA gene sequences belonging to representatives MAGs retrieved in this study are indicated in bold. Previously published[1] ASVs belonging to undefined *Ca.* Electronema are indicated in blue. The alignment used for the tree applied a 20% conservational filter to remove hypervariable positions, giving 1112 aligned positions. Boxes are used to indicate the coverage of FISH probes and the probes designed in this study are indicated in purple. Bootstrap values from 1000 re-samplings are indicated for branches with >70% (gray dot) and >90% (black) support. 16S rRNA gene sequences from isolate members of the *Desulfobulbaceae* were used as outgroup. The scale bar represents substitutions per nucleotide base.

[1] Scholz, V. V. *et al.* Cable bacteria at oxygen-releasing roots of aquatic plants: a widespread and diverse plant–microbe association. *New Phytologist* **232**, 2138–2151 (2021).

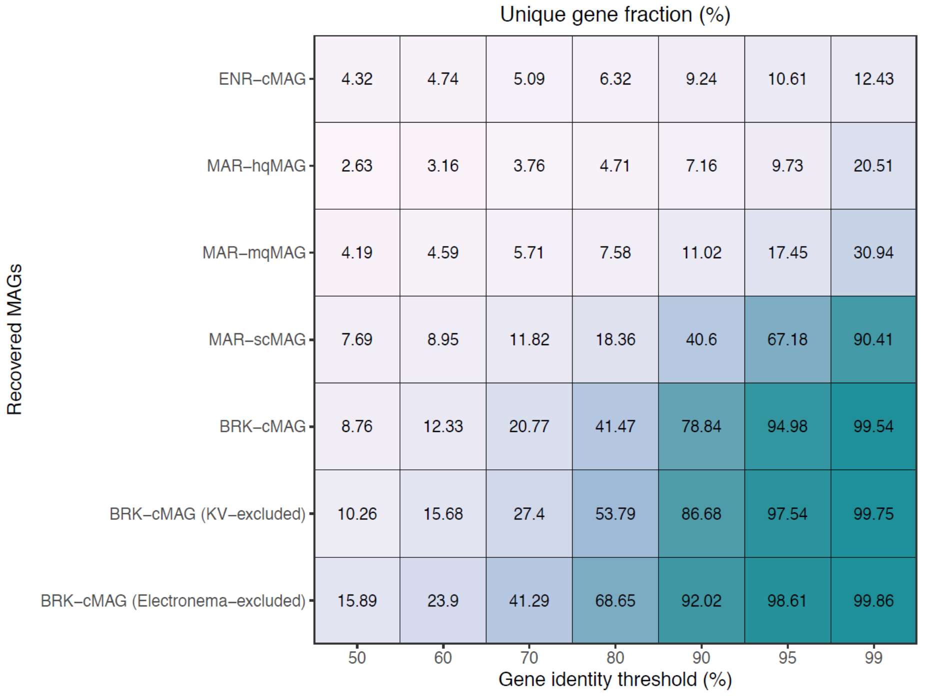


**Figure S4**. Unique gene fractions for cable bacteria MAGs recovered in this study, compared to all cable bacteria reference MAGs (HQ and MQ), at different identity thresholds for gene clustering. “-excluded” refers to the unique gene fraction that was calculated by excluding a specified MAG or MAG group from gene clustering.

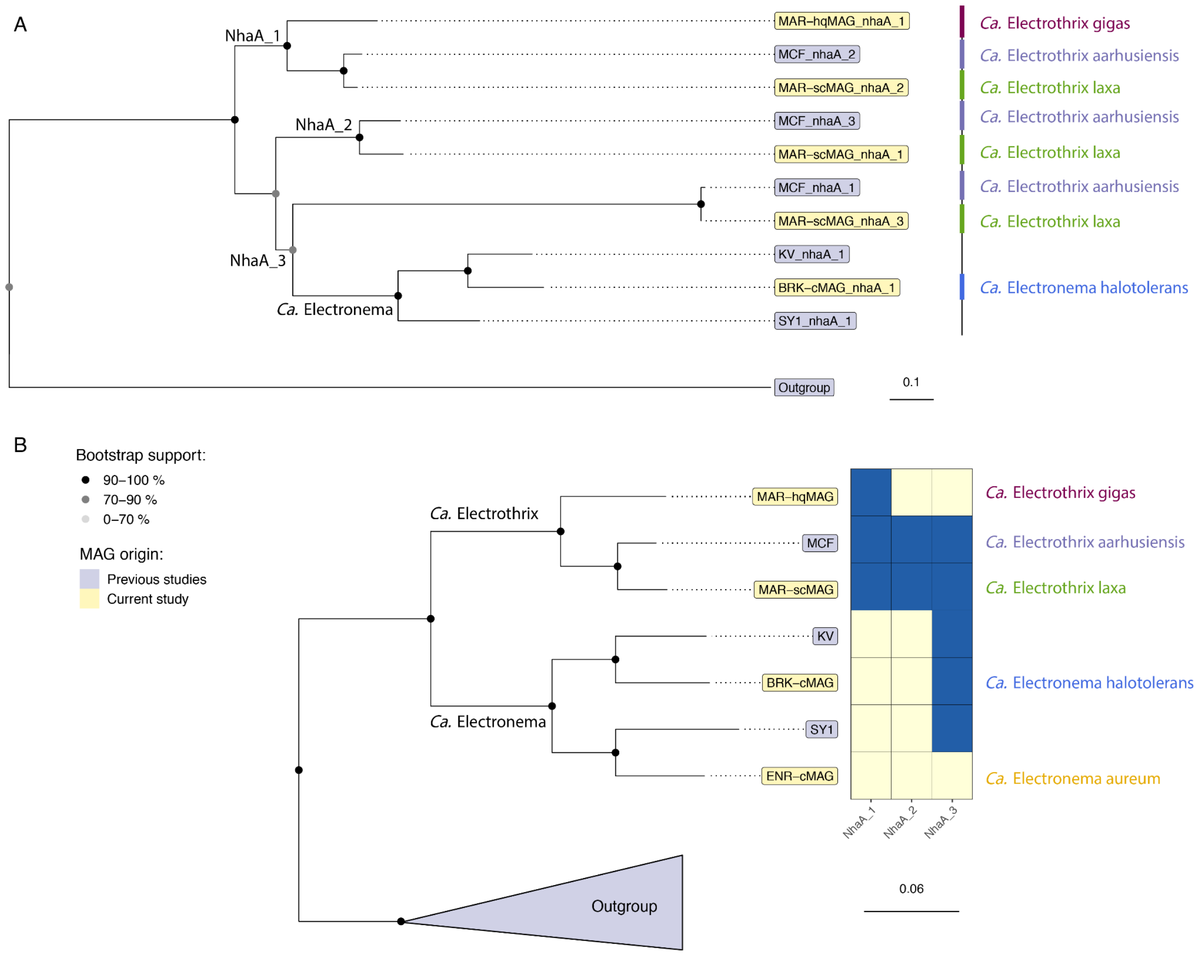


**Figure S5.** Distribution of the NhaA Na+/H+ antiporter in cable bacteria genomes. **A)** Phylogenetic tree for NhaA protein sequences, recovered from high-quality cable bacteria MAGs, which were subsetted to species representatives. Different clades of NhaA sequences have been denoted in the tree. **B)** Phylogenetic tree for cable bacteria MAGs used in **A** alongside NhaA clade presence and absence status.

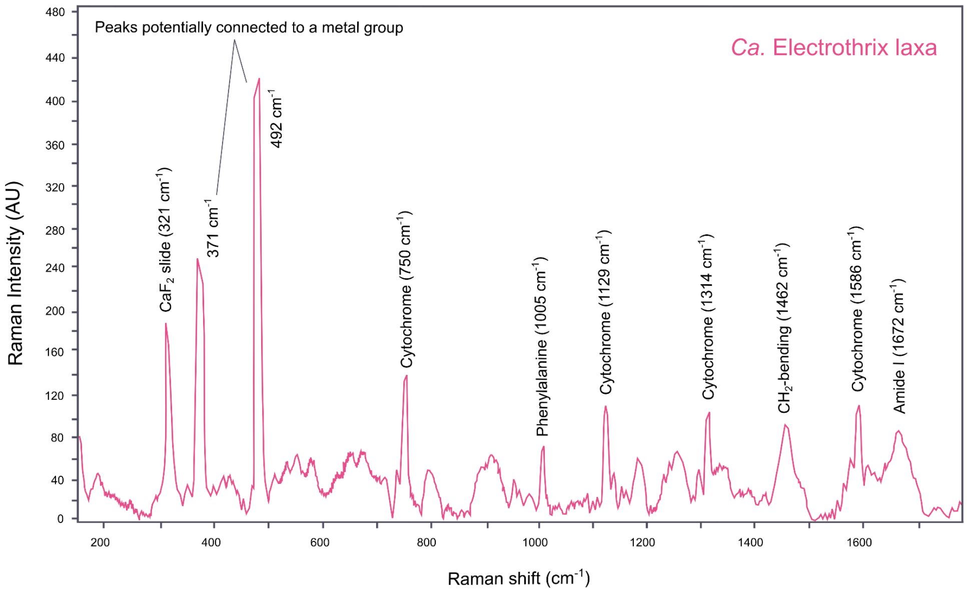


**Figure S6.** Example of a Raman spectrum from the cable bacterium *Ca*. Electrothrix laxa. The most evident peaks are highlighted in the figure: two low-frequency peaks (371 $cm^{-1}$ and 492 $cm^{-1}$) and the phenylalanine (1005 $cm^{-1}$), $CH_2$-bending (1462 $cm^{-1}$) and amide I (1672 $cm^{-1}$) peaks are indicating the presence of a potential metal group ligated to proteins in the conductive structure. Typical peaks indicating the presence of cytochromes are also present (750 $cm^{-1}$, 1129 $cm^{-1}$, 1314 $cm^{-1}$ and 1586 $cm^{-1}$).

**Table S1**. General sequencing and MAG recovery statistics for the sequenced samples. *Fraction of assembly size accounted by HQ and MQ MAGs. **For BRK and MAR samples, cable bacteria genomes were recovered after subsetting read data and performing reassembly, manual binning.

| | **ENR** | **BRK** | **MAR** |
|---|---|---|---|
| **Illumina read yield (Gb)** | 43.2 | 48.3 | 56.8 |
| **Illumina read N50 (b)** | 151 | 151 | 151 |
| **Nanopore read yield (Gb)** | 46.5 | 53.3 | 62.6 |
| **Nanopore read N50 (kb)** | 10.2 | 6.2 | 11.1 |
| **Assembly size (Mb)** | 734.6 | 2,427.5 | 1,817.5 |
| **Contigs (> 1 kb)** | 2,449 | 129,269 | 40,445 |
| **Circular contigs (> 0.5 Mb)** | 73 | 1 | 0 |
| **Contig N50 (kb)** | 3,232.1 | 38.2 | 85.4 |
| **HQ MAGs** | 58 | 32 | 13 |
| **MQ MAGs** | 8 | 115 | 72 |
| **Contigs per HQ MAG (median)** | 1 | 61 | 23 |
| **Contigs binned in HQ/MQ MAGs (%)*** | 36.7 | 23.5 | 16.1 |
| **Nanopore reads mapped to HQ/MQ MAGs (%)** | 35.5 | 35.7 | 15.5 |
| **Illumina reads mapped to HQ/MQ MAGs (%)** | 37.0 | 39.6 | 17.5 |
| **Cable bacteria MAG status**** | 1 cMAG | 1 cMAG | 2 HQ, 1 MQ |

**Table S2**. Recovered cable bacteria MAG statistics. *CDS statistics were acquired with Prokka. **Species taxonomy was determined using GTDB-tk.

| | **ENR-cMAG** | **BRK-cMAG** | **MAR-scMAG** | **MAR-hqMAG** | **MAR-mqMAG** |
|---|---|---|---|---|---|
| **MAG Completeness, %** | 93.2 | 94.1 | 92.0 | 90.2 | 93.9 |
| **MAG Contamination, %** | 3.0 | 3.3 | 2.6 | 4.0 | 7.1 |
| **Genome size, Mb** | 2.9 | 3.2 | 4.1 | 3.2 | 4 |
| **Contig count** | 1 | 1 | 1 | 37 | 9 |
| **Contig N50, Mb** | 2.9 | 3.2 | 4.1 | 0.2 | 0.9 |
| **GC rate, %** | 51.6 | 54.8 | 47.1 | 46.0 | 47.5 |
| **SNP rate, %** | 0.002 | 0.03 | 0.8 | 1.4 | 1.0 |
| **CDS count*** | 2,847 | 2,807 | 3,409 | 2,849 | 3,484 |
| **Hypothetical CDS, %*** | 56.4 | 52.3 | 58.5 | 55.5 | 60.1 |
| **16S rRNA count** | 2 | 2 | 1 | 2 | 2 |
| **23S rRNA count** | 2 | 2 | 1 | 2 | 2 |
| **5S rRNA count** | 2 | 2 | 1 | 2 | 2 |
| **MIMAG ranking** | HQ | HQ | HQ | HQ | MQ |
| **Species taxonomy ranking**** | *Ca.* Electronema GS | Novel *Ca.* Electronema | Novel *Ca.* Electrothrix | *Ca.* Electrothrix gigas | *Ca.* Electrothrix aarhusiensis |

**Table S3**. Comparison between *Ca.* Electronema GS MAGs. *Unaligned lengths determined by using the counterpart MAG as a reference sequence. **CDS predicted with Prokka automated annotation pipeline. ***Overlapping genes were determined by clustering at 70 % identity.

| | **ENR-cMAG** | **GS** |
|---|---|---|
| **MAG status** | Current study | Previous study |
| **Sequencing platform** | Nanopore long reads | Illumina short reads |
| **ID** | GCA_942492785 | GCA_004284765.1 |
| **MIMAG quality ranking** | HQ MAG | HQ MAG |
| **Genome size, Mb** | 2.93 | 2.76 |
| **Contig count** | 1 | 73 |
| **Contig N50, kb** | 2,938.1 | 58.4 |
| **16S rRNA count** | 2 | 2 |
| **Insertion sequence elements** | 163 | 38 |
| **Unaligned lengths, kb*** | 145.9 | 53.6 |
| **Unaligned contigs** | 0 | 4 |
| **Predicted CDS**** | 2，847 | 2,555 |
| **Overlapping genes***** | 2,471 | |

**Table S4**. Comparison between *Ca.* Electrothrix gigas HQ MAGs. *CDS predicted with Prokka automated annotation pipeline. **Core and unique genes were determined by clustering at 70 % identity.

| | **MAR-hqMAG** | **AW-1** | **AW-3_4** | **AW-5** | **AU-1_5** | **AS-4_5** | **AX-1_4** |
|---|---|---|---|---|---|---|---|
| **MAG status** | Current study | Previous study | | | | | |
| **Sequencing platform** | Nanopore long reads | Illumina short reads | | | | | |
| **Genome size, Mb** | 3.2 | 3.4 | 3.5 | 3.3 | 3.6 | 3.2 | 3.4 |
| **Contig count** | 37 | 407 | 354 | 444 | 163 | 808 | 320 |
| **Contig N50, kb** | 181.7 | 15.2 | 16.0 | 12.5 | 42.3 | 5.6 | 17.8 |
| **16S rRNA count** | 2 | 1 | 1 | 1 | 1 | 1 | 1 |
| **Insertion sequence elements** | 32 | 11 | 15 | 15 | 37 | 18 | 16 |
| **Predicted CDS*** | 2,849 | 3,090 | 3,129 | 3,049 | 3,109 | 2,968 | 3,092 |
| **Core genes**** | 2,150 | | | | | | |
| **Unique genes**** | 214 | 71 | 322 | 252 | 292 | 427 | 66 |

**Table S5**. Identifiers for reference genomes used in the study.

| Alias | Accession number |
|---|---|
| AW−3_4 (*Ca.* Electrothrix gigas) | GCA_022766005.1 |
| AW−5 (*Ca.* Electrothrix gigas) | GCA_022766025.1 |
| AX−2 (*Ca.* Electrothrix gigas) | GCA_022766065.1 |
| AU−1_5 (*Ca.* Electrothrix gigas) | GCA_022766165.1 |
| LOE−1_4_5 (*Ca.* Electrothrix gigas) | GCA_022766095.1 |
| AW−2 (*Ca.* Electrothrix gigas) | GCA_022765945.1 |
| AUS−3 (*Ca.* Electrothrix gigas) | GCA_022765875.1 |
| AX−1_4 (*Ca.* Electrothrix gigas) | GCA_022766045.1 |
| AW−1 (*Ca.* Electrothrix gigas) | GCA_022765985.1 |
| AS−4_5 (*Ca.* Electrothrix gigas) | GCA_022765825.1 |
| SY2 (*Ca.* Electrothrix) | GCA_011389815.1 |
| MAN−1_4 (*Ca.* Electrothrix) | GCA_022766125.1 |
| ATG−1 (*Ca.* Electrothrix) | GCA_022765905.1 |
| MCF (*Ca.* Electrothrix aarhusiensis) | GCA_004028505.1 |
| AX-5 (*Ca.* Electrothrix aarhusiensis) | GCA_022766085.1 |
| GM−3_4 (*Ca.* Electrothrix) | GCA_022765805.1 |
| A5 (*Ca.* Electrothrix marina) | GCA_004028495.1 |
| AR−5 (*Ca.* Electrothrix) | GCA_022765745.1 |
| LOE−2 (*Ca.* Electrothrix) | GCA_022766145.1 |
| EH−2 (*Ca.* Electrothrix) | GCA_022765845.1 |
| AUS4 (*Ca.* Electrothrix) | GCA_022765925.1 |
| AUS1_2 (*Ca.* Electrothrix) | GCA_022765865.1 |
| AR-3 (*Ca.* Electrothrix) | GCA_022765765.1 |
| AR-4 (*Ca.* Electrothrix) | GCA_022765725.1 |
| TB (*Ca.* Electrothrix Japonica) | SAMN03420854 |
| US4 (*Ca.* Electrothrix communis), | SAMN03420865 |
| US1 (*Ca.* Electrothrix communis) | SAMN03420863 |
| F5 (*Ca.* Electronema nielsenii) | SAMN03420862 |
| KV (*Ca.* Electronema) | PRJNA889212 |
| UQ (*Ca.* Electronema) | GCA_002413355.1 |
| GS (*Ca.* Electronema aureum) | GCA_004284765.1 |
| SY1 (*Ca.* Electronema*)* | GCA_011391865.1 |
| *Desulfobulbus alkaliphilus* | NZ_JAFFQC010000100.1 |
| *Desulfobulbus elongatus* | NZ_JHZB01000001.1 |
| *Desulfobulbus oligotrophicus* | NZ_CP054140.1 |
| *Desulfobulbus oralis* | NZ_CP021255.1 |
| *Desulfobulbus propionicus* | NC_014972.1 |
| *Desulfococcus multivorans* | NZ_CP019913.2 |
| *Desulfomonile tiedjei* | NC_018025.1 |
| *Desulfovibrio vulgaris* | NC_008751.1 |

**Table S6**. Detailed summary of the probes designed and optimized in this study.

| Probe | *E. coli* position | Specificity | Sequence (5'-3') | [FA]%** |
|---|---|---|---|---|
| **EN-logt-80** | 121-145 | *Ca. Electronema halotolerans* | 5´- CGC CAC TTT CGA TTC TCC GAA GAA -3´ | 35 |
| **EX-lin-189** | 189-210 | *Ca. Electrothrix laxa* | 5´- CCG CCT TTC TTG ATC GCC CTT -3´ | 30 |
| **EX-lin-189_C1** | 189-210 | **-** | 5´- CGC CTT TCT TGA TCG TCC TT -3´ | **-** |

**Table S7**. Overview of the data available from this study. Dereplicated HQ and MQ MAGs, which were recovered in this study and are not of cable bacteria, can be found by following the sequenced reads accession numbers at ENA.

| **Name** | **Data type** | **Accession Number** |
|---|---|---|
| ENR-NP-R941 | Sequenced reads (Nanopore R9.4.1) | ERS11891748 |
| ENR-IL | Sequenced reads (Illumina NovaSeq) | ERS11891749 |
| MAR-NP-R941 | Sequenced reads (Nanopore R9.4.1) | ERS11891750 |
| MAR-IL | Sequenced reads (Illumina NovaSeq) | ERS11891751 |
| BRK-NP-R941 | Sequenced reads (Nanopore R9.4.1) | ERS11891752 |
| BRK-IL | Sequenced reads (Illumina NovaSeq) | ERS11891753 |
| ENR-cMAG | Cable bacteria MAG (circular) | GCA_942492785 |
| BRK-cMAG | Cable bacteria MAG (circular) | GCA_942493095 |
| MAR-scMAG | Cable bacteria MAG (single-contig) | GCA_942492895 |
| MAR-hqMAG | Cable bacteria MAG (high quality) | GCA_942491745 |
| MAR-mqMAG | Cable bacteria MAG (medium quality) | GCA_942491045 |

**Table S8**. Complete protologue table for *Ca*. Electrothrix laxa.

| | |
|---|---|
| **Species name** | *Candidatus* Electrothrix laxa |
| **Genus name** | *Candidatus* Electrothrix |
| **Specific epithet** | laxa |
| **Type species of the genus** | *Candidatus* Electrothrix aarhusiensis |
| **Genus status** | Candidatus |
| **Species etymology** | Description of '*Candidatus* Electrothrix laxa' sp. nov.: "*Candidatus* Electrothrix laxa", (la´xa, L. fem. adj. *laxa*, large). This taxon is represented by the MAG MAR-scMAG. |
| **Species status** | sp. nov. |
| **Designation of the type MAG** | MAR-scMAG |
| **MAG/SAG accession number** | GCA_942492895 |
| **Genome status** | Draft |
| **Genome size** | 4075262 bp |
| **GC mol %** | 47.1 |
| **Country of origin** | Denmark |
| **Region of origin** | Central Jutland |
| **Source of sample** | Marine sediment |
| **Geographical location** | Aarhus bay |
| **Latitude** | 55°54'36.8"N |
| **Longitude** | 10°14'46.8"E |
| **Altitude/Depth** | 0 +/- 1m (due to being an intertidal area) |
| **Temperature of the sample** | 8.8–9.3C |
| **Relationship to oxygen** | Facultative anaerobe |
| **Energy metabolism** | Chemolithoautotrophs and mixotroph |
| **Assembly** | Long-read assembly, short-read polishing |
| **Sequencing technology** | Oxford Nanopore R9.4.1 and Illumina NovaSeq |
| **Binning software used** | Binning software not used |
| **Assembly software used** | Flye v. 2.9 |
| **Habitat** | Intertidal zone (coastal habitat) |
| **Miscellaneous, extraordinary features relevant for the description** | Single-contig MAG achieved after read subsetting and re-assembly |

**Table S9**. Complete protologue table for *Ca*. Electronema halotolerans.

| | |
|---|---|
| **Species name** | *Candidatus* Electronema halotolerans |
| **Genus name** | *Candidatus* Electronema |
| **Specific epithet** | halotolerans |
| **Type species of the genus** | *Candidatus* Electronema aureum |
| **Genus status** | Candidatus |
| **Species etymology** | Description of '*Candidatus* Electronema halotolerans' sp. nov.: "*Candidatus* Electronema halotolerans", (ha.lo.to´le.rans, from Gr. n. *hals*, halos salt; L. part. adj. *tolerans* tolerating; N.L. part. adj. halotolerans). This taxon is represented by the MAG BRK-cMAG. |
| **Species status** | sp. nov. |
| **Designation of the type MAG** | BRK-cMAG |
| **MAG/SAG accession number** | GCA_942493095 |
| **Genome status** | Draft |
| **Genome size** | 3199950 bp |
| **GC mol %** | 54.8 |
| **Country of origin** | Denmark |
| **Region of origin** | Central Jutland |
| **Source of sample** | Brackish sediment |
| **Geographical location** | Løgten |
| **Latitude** | 56°17'17.8"N |
| **Longitude** | 10°22'54.9"E |
| **Altitude/Depth** | 0 +/- 1m (due to being an intertidal area) |
| **Temperature of the sample** | 8.8–9.3C |
| **Relationship to oxygen** | Facultative anaerobe |
| **Energy metabolism** | Chemolithoautotrophs and mixotroph |
| **Assembly** | Long-read assembly, short-read polishing |
| **Sequencing technology** | Oxford Nanopore R9.4.1 and Illumina NovaSeq |
| **Binning software used** | Binning software not used |
| **Assembly software used** | Flye v. 2.9 |
| **Habitat** | Intertidal zone (coastal habitat) |
| **Miscellaneous, extraordinary features relevant for the description** | Circular, single-contig MAG achieved after read subsetting and re-assembly |

**Supplementary Note 1**. Additional notes on cable bacteria metabolism.

**Sulfur metabolism and electron transport.** All cable bacteria MAGs encode all key genes for the 2-step process, which includes an initial sulfide oxidation and subsequent sulfur disproportionation to sulfide and sulfate. These steps are mediated by a sulfide-quinone reductase (SQR), a polysulfide reductase (PSR) and the reversed dissimilatory sulfate reduction (DSR) pathway (**Dataset S1**). Homologs of a cytoplasmic rhodanese and the sulfur transferases TusA, likely involved in sulfur transport across the cytoplasmic membrane[1], were also encoded by all MAGs (**Dataset S1**). The resulting sulfite is oxidized to sulfate by adenosine-5-phosphosulfate reductase (AprAB) and sulfate adenylyltransferase (Sat) mediate sulfite oxidation to sulfate, which is then transported out of the cell by the sulfate permase SulP (**Dataset S1**).

Electrons released in the process are hypothesized to be transferred to the quinone pool in the cytoplasmic membrane by the heterodisulfide reductase (DsrMK) and the quinone-modifying oxidoreductase (QmoABC) membrane complexes and all genes encoding these proteins were detected in all the MAGs (**Dataset S1**). According to the proposed model for LDET, electrons are transferred to the periplasm by either a cytochrome bc-like complexe, formed by a Rieske Fe-S domain protein with an adjacent membrane-bound cytochrome b-domain protein, and/or a homolog of CydA, subunit of the membrane-bound bd quinol oxidase. The potential for both these modules was encoded in the MAGs, while no complete terminal cytochrome bd-II oxidase was detected in the genomes (**Dataset S1**). Soluble periplasmic c-type cytochromes are then hypothesized to transfer the electrons to the conductive structure of the cable bacteria, also showed experimentally with Raman microspectroscopy[2], and the genomes encode several c-type cytochromes, for instance homologs for the periplasmic cytochrome c peroxidase (Ccp), the ferrocytochrome c-552 (Cyt), or the diheme cytochrome MacA[1] (**Dataset S1**). The MAGs encode also for genes associated with type IV pili, including PilA, which could be part of the conductive fiber[1,3] (Dataset S1). The genome of *Ca.* Electrothrix laxa encodes all four subunits of the membrane-bound cytochrome c oxidase (Cco), with ~60% amino acid sequence identity to homologs in other *Desulfobulbaceae* species and most likely resulting of horizontal gene transfer, as previously observed for its close relative *Ca.* Electrothrix communis[1]. The other cable bacteria genomes did not encode membrane bound cytochrome oxidases, but homologs of the truncated hemoglobin proposed to participate in oxygen metabolism are present in all the MAGs[1] (**Dataset S1**).

Additionally, all the cable bacteria MAGs encode the NAD(P)H-quinone oxidoreductase complex (Nuo), excluding the NuoEFG subunits, and and F-type ATPase, to exploit the proton motive force (**Dataset S1**), as previously observed[1,3].

**Carbon metabolism and storage compounds**. Cable bacteria are also known for the potential to fix $CO_2$ and $CO_2$ uptake has been previously experimentally confirmed[1]. All the MAGs encode the full Wood–Ljungdahl pathway and a periplasmic carbonic anhydrase (Cah), which converts bicarbonate into $CO_2$ (**Dataset S1**). Additionally, the linked *hdrABC* genes encoding for the heterodisulfide reductase (HdrABC), potentially involved in $CO_2$ fixation[1], are encoded by *Ca*. Electrothrix laxa and *Ca*. Electronema halotolerans, and linked to a formate dehydrogenase (**Dataset S1**), and immediately followed by a homolog of *mvhD* encoding a methyl-viologen-reducing hydrogenase. Several pairs of HdrA-MvhD or HdrC-MvhD homologs are also observed in all the cable bacteria MAGs and are hypothesized to act as electron transfer modules in cytoplasmic redox reactions[1]. Additional carbon sources may be formate, which transport can be facilitated by a putative formate transporter (focA) and assimilated via the Wood–Ljungdahl pathway, or acetate, taken up by the symporter ActP and assimilated by the enzyme acetyl-CoA synthetase (Acs).

The presence of polyphosphate (poly-P) granules in cable bacteria has been proven experimentally by microscopy or nanoSIMS and they are potentially involved as protection from oxidative stress and/or as a resource to support motility[1,4]. All the cable bacteria encode the genes for polyphosphate storage, including the phosphate transporters PstABC and Pit, and the polyphosphate kinases ppk1 and ppk2 (**Dataset S1**). Furthermore, all the MAGs encode the genetic potential for production and degradation of glycogen storages, which can be used as carbon and energy resource, as well as being involved in oxygen detoxification[1].
